# Optical Control of Neuronal Activities with Photoswitchable Nanovesicles

**DOI:** 10.1101/2022.06.10.495373

**Authors:** Hejian Xiong, Kevin A. Alberto, Jonghae Youn, Jaume Taura, Johannes Morstein, Xiuying Li, Yang Wang, Dirk Trauner, Paul A. Slesinger, Steven O. Nielsen, Zhenpeng Qin

**Affiliations:** Department of Mechanical Engineering, The University of Texas at Dallas, Richardson, TX 75080, U.S.A.; Department of Chemistry, The University of Texas at Dallas, Richardson, TX 75080, U.S.A.; Nash Family Department of Neuroscience, Icahn School of Medicine at Mount Sinai, New York, NY 10029, U.S.A.; Department of Chemistry, New York University, New York, NY 10012, U.S.A.; Department of Bioengineering, The University of Texas at Dallas, Richardson, TX 75080, U.S.A.; Department of Surgery, University of Texas at Southwestern Medical Center, Dallas, TX 75080, U.S.A.; Center for Advanced Pain Studies, The University of Texas at Dallas, Richardson, TX 75080, U.S.A.

**Keywords:** azobenzene, photoswitch, liposome, controlled release, neuromodulation

## Abstract

Precise modulation of neuronal activity by neuroactive molecules is essential for understanding brain circuits and behavior. However, tools for highly controllable molecular release are lacking. Here, we developed a photoswitchable nanovesicle with azobenzene-containing phosphatidylcholine (azo-PC), coined ‘azosome’, for neuromodulation. Irradiation with 365 nm light triggers the *trans*-to-*cis* isomerization of azo-PC, resulting in a disordered lipid bilayer with decreased thickness and cargo release. Irradiation with 455 nm light induces reverse isomerization and switches the release off. Real-time fluorescence imaging shows controllable and repeatable cargo release within seconds (< 3 s). Importantly, we demonstrate that SKF-81297, a dopamine D1-receptor agonist, can be released from the azosome to activate cultures of primary striatal neurons. Azosome shows promise in precise optical control over the molecular release and can be a valuable tool for molecular neuroscience studies.

---

Understanding the role of neuroactive compounds in brain circuits and behavior is fundamentally essential in neuroscience. The delivery of neuroactive compounds via permanently implanted cannulae within the brain allows studying the effect of molecules on neural activity and behavior,^1^ but it may cause neuroinflammation and limit the activities of animals.^2^ The remotely controlled release of neuroactive compounds using external stimuli, such as a magnetic field,^3^ focused ultrasound^4, 5^ or light^6^ offers new possibilities in the area. Among these methods, light benefits from high temporal and spatial resolution. In terms of light stimulation, one of the main methods is to ‘cage’ a molecule of interest to block the activity with a photoremovable protecting group.^7, 8^ Caged compounds, such as caged glutamate or GABA, are widely used by neurophysiologists to alter cell signaling in neurons and glia, and have contributed significantly to our understanding of the nervous system.^9^ However, caged compounds have several limitations, such as challenges in ensuring biological inertness, solubility, uncaging efficiency^10^, and potential off-target activity.^11, 12^ Light-triggered release from nanocarriers is a powerful strategy for neuromodulation based on the photothermal,^13–15^ photochemical,^16–18^ or photomechanical effects.^19, 20^ However, the photorelease is not temporally controlled and cannot switch off. It remains a challenge to develop a fast and photoswitchable nanosystem that responds to a low light intensity without associated side effects (such as heating, generation of reactive oxygen species).

Azobenzene groups have been extensively used to build various bioactive photoswitches, and many have been applied in neuroscience using one-photon and two-photon stimulation.^21–28^ Recently, this approach has been expanded to lipids, and a series of photoswitchable lipids have been developed, including a photoswitchable phosphatidylcholine derivative, azo-PC. Azo-PC has enabled optical control of membrane organization and permeability.^29–33^ Chander *et al.* demonstrated light-induced release of the anticancer drug doxorubicin from azo-PC containing lipid nanoparticles *in vitro* and *in vivo*.^34^ However, it remains unclear whether azo-PC would allow the highly controllable and photoswitchable release, a critical need for neuromodulation applications.

Here we have developed a new class of photoswitchable nanovesicles, coined ‘azosomes’, by incorporating azo-PC into liposome formulations to deliver neuroactive compounds. Azo-PC undergoes reversible isomerization upon irradiation with UV-A and blue light, which rapidly changes the thickness of the lipid bilayer. The light-induced change of the lipid bilayer leads to increased permeability for encapsulated molecules and allows for switching the release of molecules on and off. Short light pulses allow temporally controlled release within seconds (< 3 s) from the photoswitchable azosome, demonstrating that the azosome is a promising tool for neuromodulation.

## Results and Discussions

### Photophysical properties of azosomes

First, we studied the reversible photoswitching property of azosomes. Figure 1A shows that one of the two lipid tails of phosphatidylcholine was modified to incorporate an azobenzene group, which can be isomerized between its *cis*- and *trans*-isomers upon the irradiation with U.V. and blue light, respectively. To increase the liposome stability, we mixed azo-PC with PEG-DSPE, DSPC, and cholesterol to prepare photoswitchable liposomes (azosomes) by the thin-film hydration method. Varying the percentage of azo-PC in the formulation leads to a series of azosomes after extruding through membranes with the pore size of 400 nm and 200 nm. The hydrodynamic diameters of these azosomes were around 160 nm (**Figure S1**). Cryo-TEM images show that most of the azosomes are spherical and unilamellar vesicles (**Figure 1B**). To investigate the photoisomerization of azo-PC in the azosome, we monitored the UV-Vis spectra of the azosome before and after the irradiation of 365 nm and 455 nm light. *Trans-azo-PC* is the thermally stable isomer, as confirmed by the strong absorption at 326 nm before light irradiation (**Figure 1C**). **Figure 1C** also shows that the absorption peak of the azosome at 326 nm decreased, and the peak at 440 nm increased in an irradiation time-dependent manner under 365 nm light. In contrast, the absorption peak at 326 nm increased, and the peak at 440 nm decreased with irradiation time under 455 nm light (**Figure 1D**). These results indicate that the irradiation of 365 nm light triggered the isomerization of *trans*-azo-PC to *cis-azo-PC.* The process was reversed under the irradiation of 455 nm light. The absolute absorbance changes at 326 nm under the irradiation of 365 nm and 455 nm overlaps well (**Figure 1E**), suggesting that the photoisomerization is reversible. The thermal *cis*-to-*trans* isomerization (relaxation) in the dark was measured at 22°C and 37°C (**Figure 1F, G, and H**). The half-lives of the relaxation were in 1-2 hours, which is much longer than the isomerization (seconds) triggered by 455 nm light. This result suggests the higher temporal control of the isomerization using 455 nm light. Next, we investigated the effect of photoisomerization of azo-PC on the azosome size. **Figure 1I** shows that the azosomes swelled by around 5 nm after the 365 nm light irradiation and can be switched back by the 455 nm light irradiation, suggesting that the reversible isomerization of azo-PC leads to reversible changes in azosome size.

**Figure 1.**
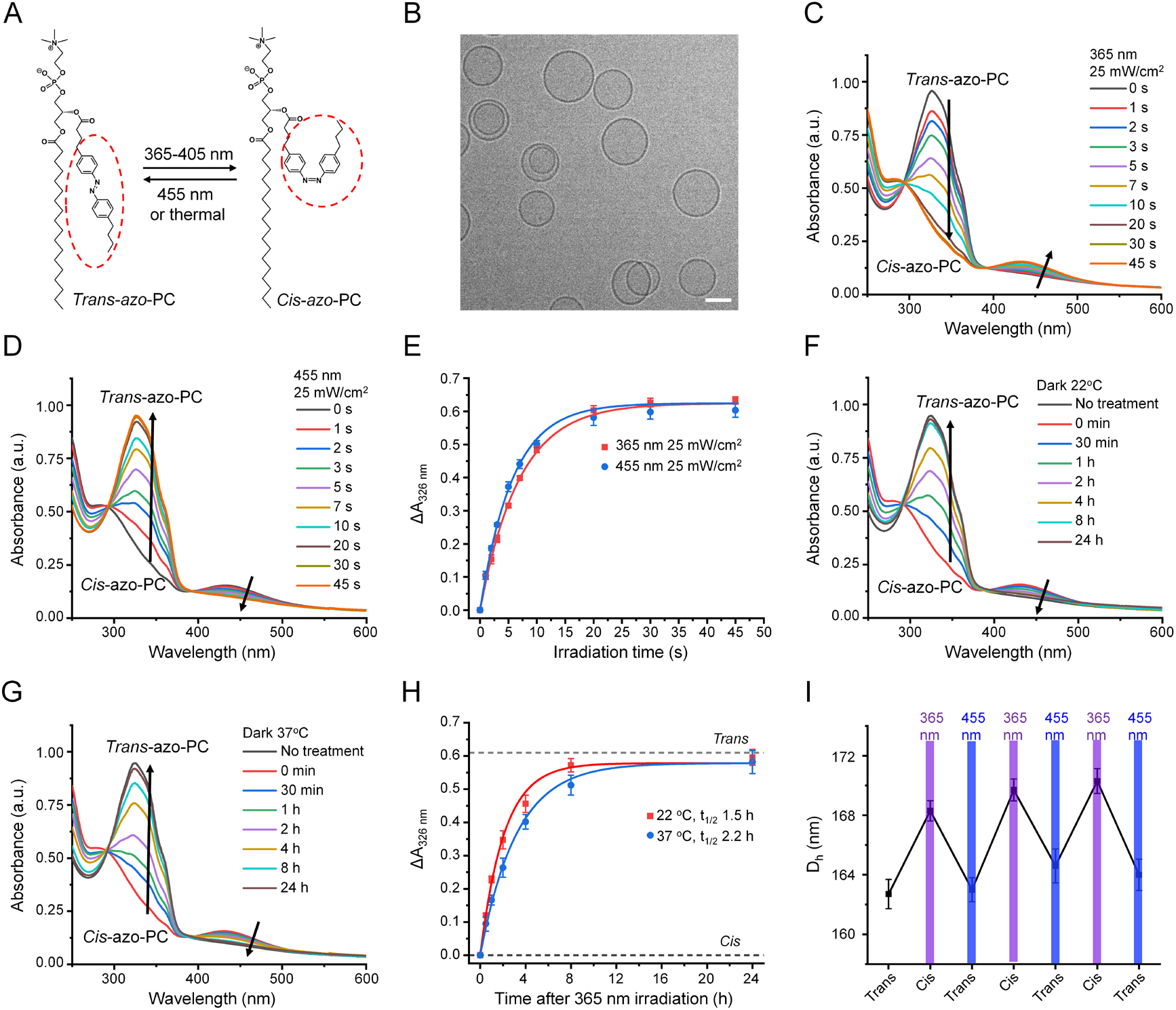
Photoswitching properties of asozomes. (A) Schematic of the photoisomerization of azo-PC. (B) Cryogenic transmission electron microscopy (Cryo-TEM) image of azosomes. Scale bar: 100 nm. (C, D) UV-Vis spectra change of azosome upon the irradiation of (C) 365 nm light and (D) 455 nm light. (E) The absorbance change of azosome at 326 nm and 455 nm light as a function of irradiation time. (F, G) UV-Vis spectra change of azosome in the dark at (F) 22°C and (G) 37°C. 365 nm light irradiation (25 mW/cm^2^, 60 s) was performed on the azosome to switch all the *trans*-azo-PC to *cis*-azo-PC at 0 min. (H) The absorbance change of azosome at 326 nm overtime in the dark at 22°C and 37°C. (I) The hydrodynamic diameter of azosome (azo-PC: 25%) under photoisomerization of azo-PC. 365 nm light or 455 nm light was irradiated on the azosome for 30 s at 25 mW/cm^2^.

### Computational modeling of azo-PC bilayer

Second, we performed all-atom molecular dynamics simulations to study the azosome bilayer at the molecular level. Simulations of pure azo-PC bilayers reveal distinct structural differences between *trans*-azo-PC and *cis*-azo-PC. **Figure 2A** shows that the azobenzene moieties align with each other in the *trans* isomer, resulting in an ordered bilayer structure. In contrast, azobenzenes in the *cis* isomer are disordered and span the bilayer hydrophobic core (**Figure 2A**). The density profiles across the bilayer confirm these differences in the molecular arrangements (**Figure 2B**). In the *cis*-azo-PC bilayer, the azo functional groups span the bilayer hydrophobic core and significantly overlap with the head group region. Notably, the bilayer thickness, defined as the peak-to-peak distance of the lipid headgroups, is smaller in the *cis*-azo-PC bilayer than the *trans*-azo-PC bilayer. We hypothesized that the disordered distribution of *cis*-azobenzene groups in the bilayer, the decreased bilayer thickness, and the lack of a methyl trough separating the upper and lower leaflets lead to the increased permeability across the *cis*-azo-PC bilayer. To demonstrate the robustness of our computational approach, we changed the torsional force field parameters of azobenzene to impose either the *cis* or *trans* state every 100 ns, resulting in reversible and reproducible bilayer properties (area per lipid shown in **Figure 2C**). These reversible bilayer changes are consistent with the experimental findings (Figure 1I). From the pure (100%) azo-PC area per lipid increase upon isomerization, we simply scaled by the azo-PC lipid composition to predict the azosome size increase as a function of azo-PC content. This linear scaling is justified by the linear nature of the experimental data (**Figure 2D**), and the comparison demonstrates that the simulation and experimental data are in reasonable agreement.

**Figure 2.**
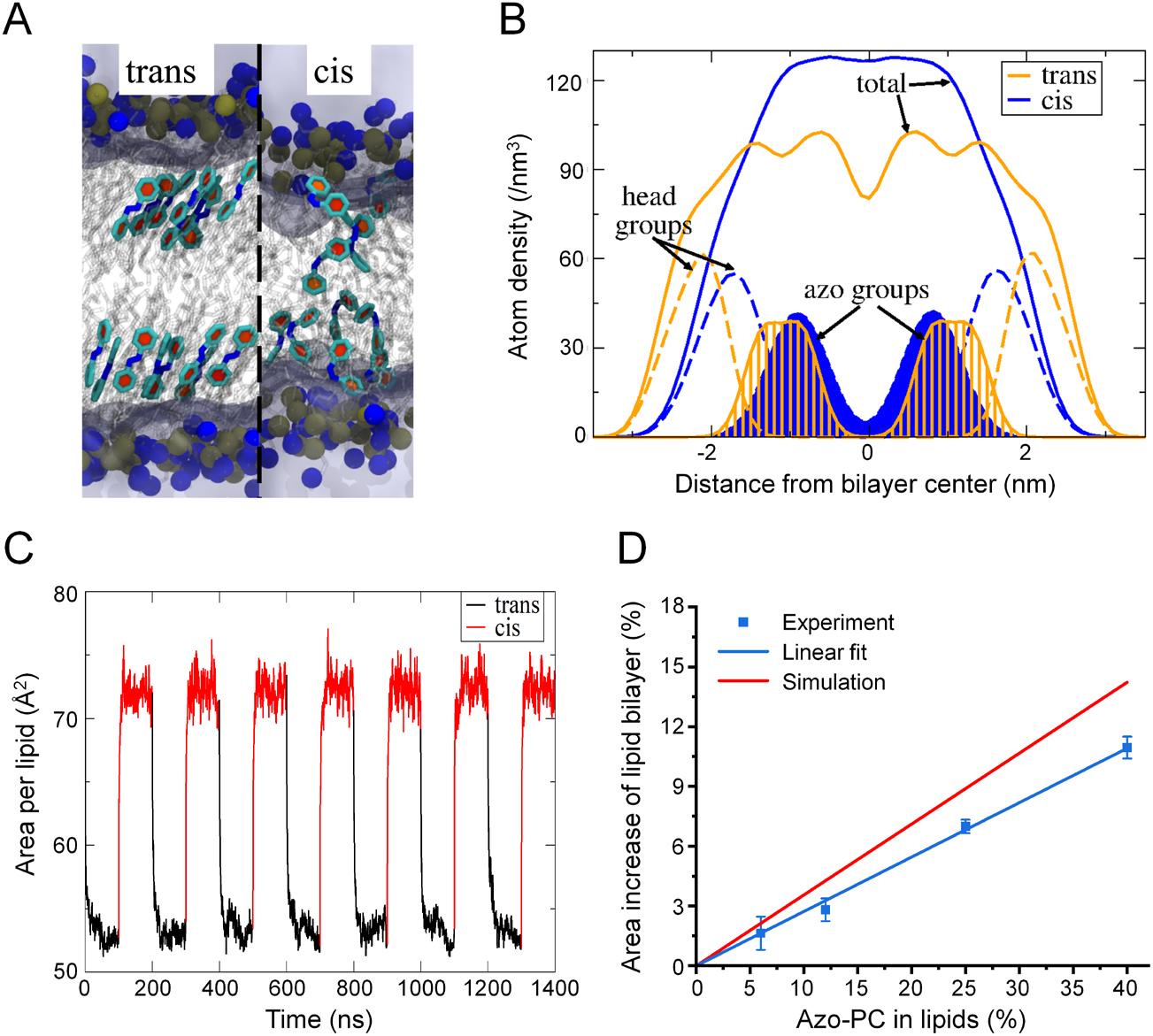
Molecular dynamics simulation for azo-PC lipid bilayer. (A) Snapshots of the *trans* and *cis* bilayers. Selected azobenzene groups are highlighted in light cyan with the ring planes in red. Lipid tails are shown in transparent gray. The blue and brown spheres represent nitrogen and phosphorous atoms on the head group, respectively. Water is represented as light purple regions. (B) The lipid atom density profile of the azo-PC bilayer as a function of the distance from the bilayer center. The orange and blue lines represent the *trans*-azo-PC and *cis*-azo-PC bilayers. Solid lines: the overall lipid density; dashed lines: lipid head groups; shaded regions: azobenzene groups. (C) Area per lipid in the tensionless ensemble of the azo-PC bilayer when the isomeric state of every lipid is altered between *cis* and *trans* every 100 ns. (D) Experimental and simulated azosome area increase with different percentages of azo-PC after the irradiation with 365 nm light (25 mW/cm^2^, 30 s). Data are expressed as Mean ± S.D. Simulation data were extrapolated to 0% azo-PC from the data in (C).

### Photoswitchable fluorophore release from azosomes

Next, we investigated the photoswitchable release from azosome by combining 365 nm and 455 nm light. We hypothesize that the *trans-to-cis* isomerization under 365 nm light irradiation increases the permeability of the liposome membrane and causes cargo release, while the reversed isomerization by 455 nm light restores the membrane integrity impeding the cargo escape from the azosome and switches off the cargo release (**Figure 3A**). To test this hypothesis, a self-quenching polyanionic dye, calcein, was encapsulated in the azosome (calcein-azosome); its fluorescence would increase upon release due to the dequenching. Minimal calcein dye leakage (< 5%) was observed with azosomes with a percentage of Azo-PC up to 25% for 24 h in 0.01 M phosphate saline buffer at room and physiological temperatures, while the 40% azo-PC formulation leads to uncontrolled leakage (**Figure S2**). This observation may be due to the low phase transition temperature with a high percentage of azo-PC. The photorelease efficiency increases with the higher azo-PC composition under 365 nm light irradiation (**Figure 3B**). We selected the calcein-azosome with 12% azo-PC for the following tests due to its high stability and photosensitivity. The photorelease efficiency is highly controllable by the intensity and duration of 365 nm and 405 nm light (**Figure 3C, Figure S3**). To explore the kinetics of the calcein release with a high temporal resolution, we recorded the real-time fluorescence of calcein using a high-speed camera. The results show that short irradiation (0.1-0.5 s) at 365 nm triggered a fast release of calcein from azosome within 1-2 s, followed by a slow-phase release that lasted 10-20 seconds (**Figure 3D, Figure S4**). To interrupt the slow-phase release, we applied second irradiation at 455 nm, which caused the stop of the release immediately (**Figure 3D**). Furthermore, the amount of released calcein is controllable by the pattern of the sequential irradiations, as shown by the higher amount of release with a 0.9 s delay between 365 nm and 455 nm light (**Figure 3D**). Interestingly, 1 s of 455 nm irradiation was sufficient to switch off the release triggered by 0.1 s of 365 nm light (**Figure 3D**), while 2 s of 455 nm irradiation was necessary to give a complete switch-off under longer 365 nm irradiation (0.2 to 0.5 s, **Figure S5**). These results suggest the ability of temporally controlled release within seconds (< 3 s). The photoswitchable properties further allow the step-wise release by multiple pulses of sequential irradiations of 365 nm and 455 nm light (**Figure 3E**). The results demonstrate that the photoswitchable azosome offers a platform for on-demand and repeatable release with both “on” and “off” switches.

**Figure 3.**
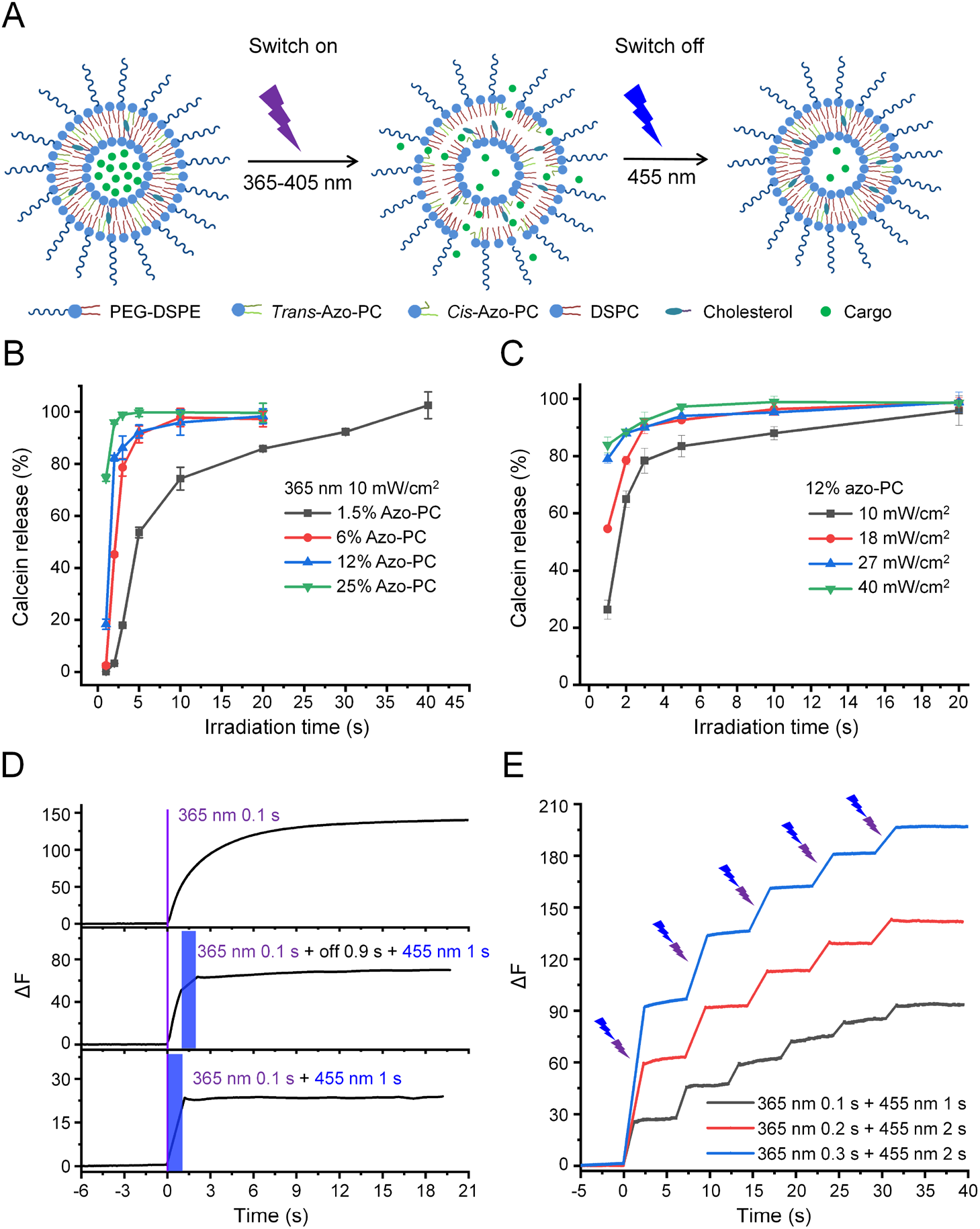
Photoswitchable fluorophore release from asozomes. (A) Schematic of the photoswitchable release from azosome. (B) Calcein release efficiency from the azosome with different percentages of azo-PC upon 365 nm light irradiation (10 mW/cm^2^). (C) Calcein release efficiency from azosome at different 365 nm light intensities and durations. Calcein fluorescence was measured 30 s after irradiation in (B) and (C). (D) Real-time fluorescence intensity of calcein over time under the sequential irradiation of 365 nm and 455 nm light (40 mW/cm^2^). The purple and blue rectangles indicate 365 nm and 455 nm light irradiation, respectively. (E) Real-time fluorescence intensity of calcein over time under multiple cycles of irradiation. The purple and blue arrows indicate the sequential irradiations of 365 nm and 455 nm light every 5 s (40 mW/cm^2^). Data are expressed as Mean ± S.D.

### Photoswitchable neuromodulator release from azosomes

Lastly, we tested the application of azosome for neuromodulation. Here we encapsulated SKF-81297, a dopamine D1-like receptor agonist, in the azosome (SKF-azosome, **Figure 4A**). Approximately half of the striatal neuron population expresses the D1 receptor.^35^ We predicted that the release of SKF-81297 would activate the D1 receptor on striatal neurons and increase intracellular Ca^2+^ and neuronal excitability.^36, 37^ Primary striatal neurons were cultured on glass-bottom dishes and used for Ca^2+^ imaging after the neurons had matured (14 days). We first stained neurons with Ca^2+^-sensitive dye Fluo-4 as a proxy for neural activity and applied Dil-labeled azosomes to the extracellular medium. After 1 h, there was no evidence of significant endocytosis of azosome by neurons, suggesting that azosomes remain predominantly extracellular (**Figure S5**). After the sequential irradiations at 365 nm and 455 nm, 56% of the neurons incubated with SKF-azosomes exhibited an increase in fluorescence, indicating an increase in intracellular Ca^2+^ (as identified by ΔF/F_0_ ≥20%, **Figure 4B**, **C**). For control, we examined the effect of irradiating SKF-liposomes lacking azo-PC and observed no change in fluorescence (**Figure 4D**). We also investigated any potential cytotoxicity; a live/dead cell viability assay indicated no significant toxicity of azosomes or the irradiation paradigm (**Figure S6**). These results demonstrate that azosome is a useful platform for the controlled release of neurochemicals for neuromodulation.

**Figure 4.**
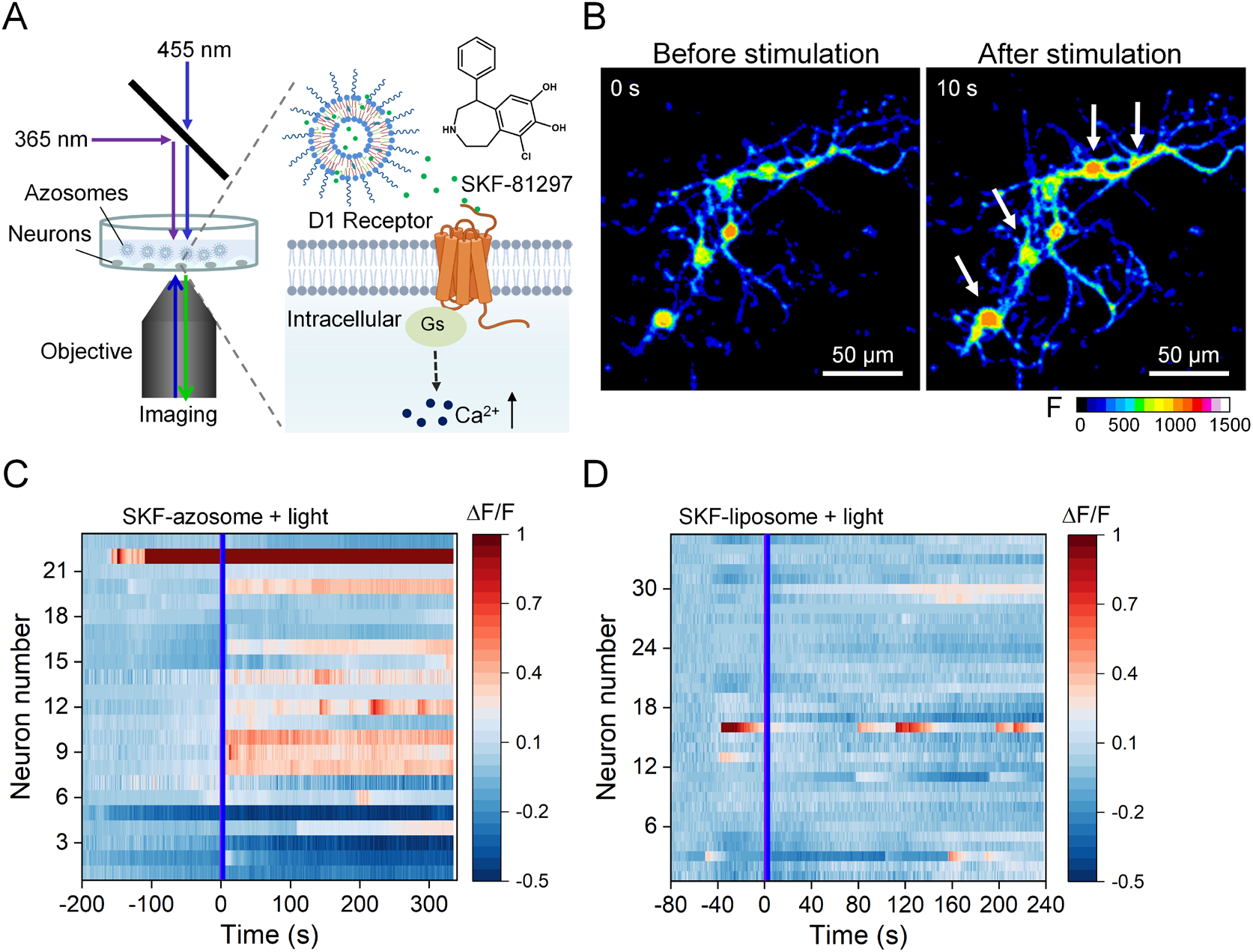
Controlled release for neuromodulation. (A) Schematic photorelease of SKF-81297 from the azosome to induce Ca^2+^ influx in primary mouse striatal neurons. (B) Real-time fluorescent image of primary mouse striatal neurons before and after the irradiation of sequential 365 nm light (40 mW/cm^2^, 0.5 s) and 455 nm light (40 mW/cm^2^, 2 s). Fluo-4 was used as the Ca^2+^ indicator. Scale bar: 50 μm. (C, D) The fluorescence change from individual neurons incubated with (C) SKF-azosome and (D) SKF-liposome. The irradiation of 365 nm and 455 nm light was performed during the imaging.

## Conclusion

In summary, we developed a photoswitchable nanovesicle, referred to as ‘azosome’, based on the reversible photoisomerization of azo-PC under the irradiation of 365 nm and 455 nm light. The *trans-to-cis* isomerization triggered by 365 nm light decreases the thickness of the liposome bilayer, increases the azosome diameter, and leads to efficient cargo release. In contrast, the reversed *cis-to-trans* isomerization induced by 455 nm light increases the thickness of the liposome bilayer, decreases the azosome diameter, and switches off the release. We demonstrate that photorelease is controllable within seconds (< 3 s), is repeatable, and can specifically activate targeted neurons by releasing the dopamine D1-like receptor agonist SKF-81297. With the emerging red-azo-PC, the azosome is expected to be photoswitched under the irradiationof tissue-penetrating red light (≥630 nm). The azosome formulation shows promise in precise optical control over the release of neuroactive compounds for neuromodulation and can be a valuable tool for molecular neuroscience studies.

## Materials and Methods

### Materials

1,2-Distearoyl-sn-glycero-3-phosphocholine (DSPC, >99%), 1,2-distearoyl-sn-glycero-3-phosphoethanolamine-N-[amino(polyethylene glycol)-2000] (ammonium salt) (DSPE-PEG2000, >99%) and cholesterol (ovine wool, >98%) were purchased from Avanti Polar Lipids, Inc. Calcein sodium salt (644.5 g/mol, 108750-13-6) was purchased from Alfa Aesar. SKF-81297 hydrobromide (370.67 g/mol, ≥ 98%) was purchased from Tocris Bioscience. 1,1’-Dioctadecyl-3,3,3’,3’-tetramethylindocarbocyanine perchlorate (DilC18(3)), ≥ 90%) was purchased from ThermoFisher Scientific. All other chemicals were analytical grade.

### Preparation and characterization of Azosome

Preparation of different azo-PC-containing liposomes was performed by the thin-film hydration method reported previously.^19, 20^ DSPC, PEG-DSPE, cholesterol, and azo-PC were mixed in chloroform at the molar ratio of 57: 1: 30: 12. The lipid film was prepared with N2 for several minutes and dried under vacuum overnight. 10 mM phosphate-buffered saline (PBS) containing either calcein (75 mM) or sulfate ammonium (300 mM) was added to the film for 1 h at 65 °C. After 5 freeze-thaw cycles (1 minute in liquid N2 and 2 minutes in 65 °C water bath, respectively), the liposome solution was extruded through 400 nm and 200 nm polycarbonate membranes (Whatman, USA) for 11 passages each using a Mini Extruder (Avanti Polar Lipids, USA). Afterward, free calcein or sulfate ammonium was removed by size exclusion chromatography with Sephacryl^®^ S-500 HR column (Cat. # GE17-0613-10, Sigma-Aldrich). The calcein-loaded azosome was used directly. To load the SKF-81297, the sulfate ammonium-loaded azosome was incubated with 1 mM SKF-81297 at 70°C for 30 min. SKF-81287 loaded azosome was further purified by dialysis. The molar percentage of azo-PC in the azosome was adjusted to 1.5%, 6%, 12%, 25%, and 40% to investigate its effect on release efficiency, and the percentage of DSPC was changed accordingly. As a control group, SKF-81297 loaded liposome (without azo-PC) was prepared by similar methods. 0.5% of DilC18(3) was mixed with lipids to label azosome (Dil-azosome).

The hydrodynamic size, size distribution, and zeta-potential of the liposomes were determined by dynamic light scattering measurement (Malvern Zetasizer Nano ZS) at room temperature. DU800 spectrophotometer (Beckman Coulter) was used to track the UV-Vis absorption spectrum change of azo-PC before and after the photoisomerization. The morphology of liposomes was observed by a transmission electron microscope with an FEI Tecnai 300kV field emission gun and a Gatan Summit K2 direct electron detector camera at the Characterization Facility, University of Minnesota.

### *In vitro* photoisomerization and release

To investigate the photoisomerization of azo-PC, 100 μL of empty azosome was added to a 96-well plate and irradiated by 365 nm and 455 nm light (25 mW/cm^2^). Afterward, the UV-Vis absorption spectra of azosome before and after irradiation were recorded immediately. To investigate the release efficiency of azosome, calcein-loaded azosome was added to a 96-well plate and irradiated by 365 nm and 455 nm light at a series of intensities or durations. The fluorescence intensity (485 nm excitation and 520 nm emission) was measured using a microplate reader (Synergy H2). For 100% release, azosomes were treated with 0.5% Triton X-100. The percentage release of calcein was calculated by the following formula:

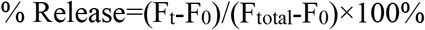

where F_0_, F_t_, and F_total_ represent the initial fluorescence signal, fluorescence signal after light irradiation, and after being treated by Triton, respectively.

The release kinetics of azosome was investigated following our previously reported method.^20^ Briefly, an aliquot of well-dispersed calcein-loaded azosome was placed on a glass slide, covered by a cover slide, and sealed by nail polish. The samples were then immobilized onto a microscope (Olympus IX73) stage and irradiated by 365 nm and 455 nm light. A high-speed digital camera (Hamamatsu Photonics, ORCA-Flash 4.0) was used to record the real-time fluorescent intensity profile every 50 ms. A series of fluorescent images were obtained, and the fluorescence intensity was analyzed by Image J.

### Primary neuron culture and neuromodulation

Cryopreserved primary mouse neurons derived from the striatum of day 18 embryonic CD1 mouse brain (Cat. # M8812N-10) and the culture kit (Cat. # M8812NK-10) were purchased from Cell Applications, Inc. The glass-bottom dish (diameter of glass: 10 mm) was treated by neuron coating solution overnight and rinsed with PBS before seeding. After thawing, the cryopreserved primary mouse neurons were transferred to a 50 mL tube containing a 4 mL mouse neuron plating medium. The neurons were then aliquoted into 8 dishes (only on the glass) to ensure the seeding density is 100, 000 cells per cm^2^ or above, and cultured in a cell culture incubator (37°C, 5% CO_2_, humidified). Half of the mouse neuron culture medium was changed every three days. After 2 weeks, the neurons were incubated with fluo-4 AM (Cat. # F14201, Molecular Probes^TM^) at the concentration of 3 μM for 30 min, and then washed with PBS three times. 300 uL of artificial cerebrospinal fluid (ACSF, 124 mM NaCl, 5 mM KCl, 26 mM NaHCO3, 1.25 mM NaH2PO4, 10 mM D-Glucose, 1.3 mM MgCl2 and 1.5 mM CaCl2) containing SKF-81297 loaded azosome (12% azo-PC, total lipids: 2 mM) was added into the glass area in the dish. The fluorescence of fluo-4 in the neurons was recorded by a spinning disk microscope (Olympus, SD-OSR) at the excitation of 488 nm. During the recording, the whole glass area was irradiated by sequential 365 nm light (0.5 s, 40 mW/cm^2^) and 455 nm light (2 s, 40 mW/cm^2^) to control the release of SKF-81297. The fluorescence intensity change (ΔF/F0) from individual neurons was analyzed by Image J. F0 value of each neuron was measured by averaging its fluorescent intensity during the 100 s before the irradiation. To investigate the distribution of azosome, Dil-azosome (total lipids: 20 mM) was incubated with neurons for 1 h and imaged by the microscope at the excitation of 561 nm. After wash, the neurons were imaged again at the same condition.

### *In vitro* toxicity

The cytotoxicity of azosomes on neurons was evaluated by the live/dead staining method. The primary striatal neurons were cultured in the 96 wells plate. After 2 weeks, the neurons were incubated with azosome (12% azo-PC, total lipids: 2 mM) for 1 h and 24 h. The light irradiation paradigm used in the neuromodulation experiments was treated on the neurons. After the treatment, the live and dead cells were stained with 3 μM calcein-AM (ThermoFisher, Cat. # C3100MP) and 5 μM propidium iodide (PI, ThermoFisher, Cat. # P1304MP) for 20 min, respectively. After wash, the fluorescence of calcein and PI was measured by the plate reader. The fluorescent images of stained cells were taken by the confocal microscope (Olympus, FV3000RS).

### All-atom modeling of the lipid bilayer

All-atom simulations of azo-PC were performed using the CHARMM36 lipid^38^ force field and the NAMD simulation software (version 2.13).^39^ The initial azo-PC structure was created in Avogadro (version 1.2.0)^40^ and minimized using the built-in steepest descent algorithm. The topology entry for azo-PC was created from the existing entry of DSPC and manually altered to incorporate the azobenzene moiety. Atomistic force field parameters of azo-PC were created from a mixture of existing DSPC parameters, previously published parameters for the azobenzene moiety,^41^ and any missing parameters were obtained through CGenFF.^42–46^ Packmol was used to create the initial bilayer structures,^47^ and the VMD Solvate Plugin was used to solvate the bilayer with the TIP3P water model.^48^ All simulations were performed in the tensionless ensemble at 1 atm and 323 K. Analysis was performed using custom Python scripts utilizing the MD Analysis package.^49, 50^

Torsional force field parameters to control the azobenzene isomeric state are used following our previous method.^41^ To mimic photoisomerization in a simulation, the torsional parameters corresponding to the desired photoisomeric state are enabled while simultaneously disabling the torsional parameters for the opposite state. This parameter change is made between simulation runs, using a restart or checkpoint file to continue with the new parameter set. Thus, for a given simulation run the parameters for only a single photoisomeric state are enabled at a time.

The following equation was used to calculate the change of the azosome surface area during isomerization from the experimental diameter data.

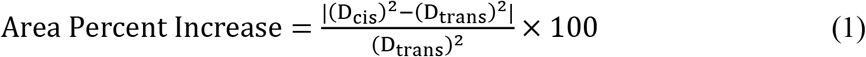

where D*_cis_* and D*_trans_* are the azosome diameters for *azo-PC* in the *cis* and *trans* state, respectively.

The areas per lipid (APL) from the simulations were calculated using the following equation:

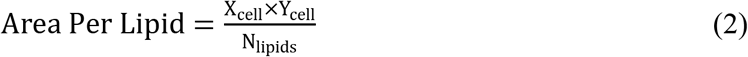

where X_cell_ and Y_cell_ are the cell dimensions in the X- and Y-directions, respectively, and N_lipids_ is the number of lipids per bilayer leaflet.

The change of bilayer area in the simulation was calculated by the following equation:

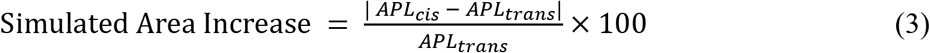

## ASSOCIATED CONTENT

### Supporting Information

The following files are available free of charge. All the experimental materials and methods, including the preparation and characterization of azosome, in vitro photoisomerization and release, primary neuron culture and neuromodulation, in vitro toxicity, all-atom modeling of the lipid bilayer; additional results Figure S1-S6 (PDF)

## AUTHOR INFORMATION

### Author Contributions

H.X., P.A.S., and Z.Q. conceived and designed the experiments. H.X., J.Y., and X.L. performed the experiments and analyzed the data. J.M. and D.T. synthesized the azo-PC. K.A.A. and Y.W. performed the theoretical modeling for lipid bilayer under the supervision of S.O.N.. J. T. assisted in the neuromodulation experiments. H.X. wrote and revised the manuscript. P.A.S., S.O.N., and Z.Q. supervised the project. All authors discussed the results and revised the manuscript.

### Notes

The authors declare that they have no competing financial interests.

## ACKNOWLEDGMENT

This work was partially supported by National Science Foundation under award number 2123971 (Z.Q., P.S., S.N.), National Institute of Neurological Disorders and Stroke of the National Institutes of Health under award number RF1NS110499 (Z.Q., P.S.), and a postdoc research grant from the Phospholipid Research Center (Heidelberg, Germany) to H.X.

## REFERENCES

1. Dagdeviren, C.; Ramadi, K. B.; Joe, P.; Spencer, K.; Schwerdt, H. N.; Shimazu, H.; Delcasso, S.; Amemori, K.-i.; Nunez-Lopez, C.; Graybiel, A. M., Miniaturized neural system for chronic, local intracerebral drug delivery. Sci. Transl. Med. 2018, 10, eaan2742.

2. Feiner, R.; Dvir, T., Tissue–electronics interfaces: from implantable devices to engineered tissues. Nat. Rev. Mater. 2017, 3, 1–16.

3. Rao, S.; Chen, R.; LaRocca, A. A.; Christiansen, M. G.; Senko, A. W.; Shi, C. H.; Chiang, P.-H.; Varnavides, G.; Xue, J.; Zhou, Y., Remotely controlled chemomagnetic modulation of targeted neural circuits. Nat. Nanotechnol. 2019, 14, 967–973.

4. Airan, R. D.; Meyer, R. A.; Ellens, N. P.; Rhodes, K. R.; Farahani, K.; Pomper, M. G.; Kadam, S. D.; Green, J. J., Noninvasive targeted transcranial neuromodulation via focused ultrasound gated drug release from nanoemulsions. Nano Lett. 2017, 17, 652–659.

5. Wang, J. B.; Aryal, M.; Zhong, Q.; Vyas, D. B.; Airan, R. D., Noninvasive ultrasonic drug uncaging maps whole-brain functional networks. Neuron 2018, 100, 728–738. e7.

6. Rapp, T. L.; DeForest, C. A., Targeting drug delivery with light: A highly focused approach. Adv. Drug Deliv. Rev. 2021, 171, 94–107.

7. Ellis-Davies, G. C., Caged compounds: photorelease technology for control of cellular chemistry and physiology. Nat. Methods 2007, 4, 619–28.

8. Taura, J.; Nolen, E. G.; Cabré, G.; Hernando, J.; Squarcialupi, L.; López-Cano, M.; Jacobson, K. A.; Fernández-Dueñas, V.; Ciruela, F., Remote control of movement disorders using a photoactive adenosine A2A receptor antagonist. J. Control. Release 2018, 283, 135–142.

9. Ellis-Davies, G. C., Useful caged compounds for cell physiology. Acc. Chem. Res. 2020, 53, 1593–1604.

10. Silva, J. M.; Silva, E.; Reis, R. L., Light-triggered release of photocaged therapeutics - Where are we now? J. Control. Release 2019, 298, 154–176.

11. Maier, W.; Corrie, J. E. T.; Papageorgiou, G.; Laube, B.; Grewer, C., Comparative analysis of inhibitory effects of caged ligands for the NMDA receptor. J. Neurosci. Methods 2005, 142, 1–9.

12. Noguchi, J.; Nagaoka, A.; Watanabe, S.; Ellis-Davies, G. C.; Kitamura, K.; Kano, M.; Matsuzaki, M.; Kasai, H., In vivo two-photon uncaging of glutamate revealing the structure–function relationships of dendritic spines in the neocortex of adult mice. J. Physiol. 2011, 589, 2447–2457.

13. Li, B.; Wang, Y.; Gao, D.; Ren, S.; Li, L.; Li, N.; An, H.; Zhu, T.; Yang, Y.; Zhang, H.; Xing, C., Photothermal modulation of depression-related ion channel function through conjugated polymer nanoparticles. Adv. Funct. Mater. 2021, 31, 2010757.

14. Nakano, T.; Mackay, S. M.; Wui Tan, E.; Dani, K. M.; Wickens, J., Interfacing with neural activity via femtosecond laser stimulation of drug-encapsulating liposomal nanostructures. eNeuro 2016, 3.

15. Li, W.; Luo, R.; Lin, X.; Jadhav, A. D.; Zhang, Z.; Yan, L.; Chan, C.-Y.; Chen, X.; He, J.; Chen, C.-H.; Shi, P., Remote modulation of neural activities via near-infrared triggered release of biomolecules. Biomaterials 2015, 65, 76–85.

16. Huu, V. A. N.; Luo, J.; Zhu, J.; Zhu, J.; Patel, S.; Boone, A.; Mahmoud, E.; McFearin, C.; Olejniczak, J.; de Gracia Lux, C.; Lux, J.; Fomina, N.; Huynh, M.; Zhang, K.; Almutairi, A., Light-responsive nanoparticle depot to control release of a small molecule angiogenesis inhibitor in the posterior segment of the eye. J. Control. Release 2015, 200, 71–77.

17. Kohman, R. E.; Cha, S. S.; Man, H. Y.; Han, X., Light-triggered release of bioactive molecules from DNA nanostructures. Nano Lett. 2016, 16, 2781–5.

18. Veetil, A. T.; Chakraborty, K.; Xiao, K.; Minter, M. R.; Sisodia, S. S.; Krishnan, Y., Cell-targetable DNA nanocapsules for spatiotemporal release of caged bioactive small molecules. Nat. Nanotechnol. 2017, 12, 1183–1189.

19. Xiong, H.; Li, X.; Kang, P.; Perish, J.; Neuhaus, F.; Ploski, J. E.; Kroener, S.; Ogunyankin, M. O.; Shin, J. E.; Zasadzinski, J. A.; Wang H.; Slesinger P. A.; Zumbuehl A.; Qin Z., Near-infrared light triggered-release in deep brain regions using ultra-photosensitive nanovesicles. Angew. Chem. 2020, 132, 8686–8693.

20. Li, X.; Che, Z.; Mazhar, K.; Price, T. J.; Qin, Z., Ultrafast near-infrared light-triggered intracellular uncaging to probe cell signaling. Adv. Funct. Mater. 2017, 27, 1605778.

21. Cabré, G.; Garrido-Charles, A.; Moreno, M.; Bosch, M.; Porta-de-la-Riva, M.; Krieg, M.; Gascón-Moya, M.; Camarero, N.; Gelabert, R.; Lluch, J. M.; Busqué, F.; Hernando, J.; Gorostiza, P.; Alibés, R., Rationally designed azobenzene photoswitches for efficient two-photon neuronal excitation. Nat. Commun. 2019, 10, 907.

22. DiFrancesco, M. L.; Lodola, F.; Colombo, E.; Maragliano, L.; Bramini, M.; Paternò, G. M.; Baldelli, P.; Serra, M. D.; Lunelli, L.; Marchioretto, M.; Grasselli, G.; Cimò, S.; Colella, L.; Fazzi, D.; Ortica, F.; Vurro, V.; Eleftheriou, C. G.; Shmal, D.; Maya-Vetencourt, J. F.; Bertarelli, C.; Lanzani, G.; Benfenati, F., Neuronal firing modulation by a membrane-targeted photoswitch. Nat. Nanotechnol. 2020, 15, 296–306.

23. Kellner, S.; Berlin, S., Two-photon excitation of azobenzene photoswitches for synthetic optogenetics. Appl. Sci. 2020, 10.

24. Morstein, J.; Dacheux, M. A.; Norman, D. D.; Shemet, A.; Donthamsetti, P. C.; Citir, M.; Frank, J. A.; Schultz, C.; Isacoff, E. Y.; Parrill, A. L.; Tigyi, G. J.; Trauner, D., Optical control of lysophosphatidic acid signaling. J. Am. Chem. Soc. 2020, 142, 10612–10616.

25. Morstein, J.; Hill, R. Z.; Novak, A. J. E.; Feng, S.; Norman, D. D.; Donthamsetti, P. C.; Frank, J. A.; Harayama, T.; Williams, B. M.; Parrill, A. L.; Tigyi, G. J.; Riezman, H.; Isacoff, E. Y.; Bautista, D. M.; Trauner, D., Optical control of sphingosine-1-phosphate formation and function. Nat. Chem. Biol. 2019, 15, 623–631.

26. Morstein, J.; Romano, G.; Hetzler, B. E.; Plante, A.; Haake, C.; Levitz, J.; Trauner, D., Photoswitchable serotonins for optical control of the 5-HT2A receptor. Angew. Chem. Int. Ed. 2022, 61, e202117094.

27. Mukhopadhyay, T. K.; Morstein, J.; Trauner, D., Photopharmacological control of cell signaling with photoswitchable lipids. Curr. Opin. Pharmacol. 2022, 63, 102202.

28. Bahamonde, M. I.; Taura, J.; Paoletta, S.; Gakh, A. A.; Chakraborty, S.; Hernando, J.; Fernández-Dueñas, V.; Jacobson, K. A.; Gorostiza, P.; Ciruela, F., Photomodulation of G protein-coupled adenosine receptors by a novel light-switchable ligand. Bioconjug. Chem. 2014, 25, 1847–1854.

29. Pernpeintner, C.; Frank, J. A.; Urban, P.; Roeske, C. R.; Pritzl, S. D.; Trauner, D.; Lohmüller, T., Light-controlled membrane mechanics and shape transitions of photoswitchable lipid vesicles. Langmuir 2017, 33, 4083–4089.

30. Pritzl, S. D.; Konrad, D. B.; Ober, M. F.; Richter, A. F.; Frank, J. A.; Nickel, B.; Trauner, D.; Lohmüller, T., Optical membrane control with red light enabled by red-shifted photolipids. Langmuir 2021, 38, 385–393.

31. Pritzl, S. D.; Urban, P.; Prasselsperger, A.; Konrad, D. B.; Frank, J. A.; Trauner, D.; Lohmüller, T., Photolipid bilayer permeability is controlled by transient pore formation. Langmuir 2020, 36, 13509–13515.

32. Urban, P.; Pritzl, S. D.; Konrad, D. B.; Frank, J. A.; Pernpeintner, C.; Roeske, C. R.; Trauner, D.; Lohmüller, T., Light-controlled lipid interaction and membrane organization in photolipid bilayer vesicles. Langmuir 2018, 34, 13368–13374.

33. Urban, P.; Pritzl, S. D.; Ober, M. F.; Dirscherl, C. F.; Pernpeintner, C.; Konrad, D. B.; Frank, J. A.; Trauner, D.; Nickel, B.; Lohmueller, T., A lipid photoswitch controls fluidity in supported bilayer membranes. Langmuir 2020, 36, 2629–2634.

34. Chander, N.; Morstein, J.; Bolten, J. S.; Shemet, A.; Cullis, P. R.; Trauner, D.; Witzigmann, D., Optimized photoactivatable lipid nanoparticles enable red light triggered drug release. Small 2021, 2008198.

35. Gagnon, D.; Petryszyn, S.; Sanchez, M.; Bories, C.; Beaulieu, J.; De Koninck, Y.; Parent, A.; Parent, M., Striatal neurons expressing D1 and D2 receptors are morphologically distinct and differently affected by dopamine denervation in mice. Sci. Rep. 2017, 7, 1–16.

36. Dai, R.; Ali, M. K.; Lezcano, N.; Bergson, C., A crucial role for cAMP and protein kinase a in D1 dopamine receptor regulated intracellular calcium transients. NeuroSignals 2008, 16, 112–123.

37. Jeroen Vermeulen, R.; Drukarch, B.; Rob Sahadat, M. C.; Goosen, C.; Wolters, E. C.; Stoof, J. C., The selective dopamine D1 receptor agonist, SKF 81297, stimulates motor behaviour of MPTP-lesioned monkeys. Eur. J. Pharmacol. 1993, 235, 143–147.

38. Klauda, J. B.; Venable, R. M.; Freites, J. A.; O’Connor, J. W.; Tobias, D. J.; Mondragon-Ramirez, C.; Vorobyov, I.; MacKerell Jr, A. D.; Pastor, R. W., Update of the CHARMM all-atom additive force field for lipids: validation on six lipid types. J. Phys. Chem. B 2010, 114, 7830–7843.

39. Phillips, J. C.; Hardy, D. J.; Maia, J. D.; Stone, J. E.; Ribeiro, J. V.; Bernardi, R. C.; Buch, R.; Fiorin, G.; Hénin, J.; Jiang, W., Scalable molecular dynamics on CPU and GPU architectures with NAMD. J. Chem. Phys. 2020, 153, 044130.

40. Hanwell, M. D.; Curtis, D. E.; Lonie, D. C.; Vandermeersch, T.; Zurek, E.; Hutchison, G. R., Avogadro: an advanced semantic chemical editor, visualization, and analysis platform. J. Cheminform. 2012, 4, 1–17.

41. Siriwardane, D. A.; Kulikov, O.; Batchelor, B. L.; Liu, Z.; Cue, J. M.; Nielsen, S. O.; Novak, B. M., UV-and thermo-controllable azobenzene-decorated polycarbodiimide molecular springs. Macromolecules 2018, 51, 3722–3730.

42. Gutiérrez, I. S.; Lin, F.-Y.; Vanommeslaeghe, K.; Lemkul, J. A.; Armacost, K. A.; Brooks III, C. L.; MacKerell Jr, A. D., Parametrization of halogen bonds in the CHARMM general force field: Improved treatment of ligand–protein interactions. Bioorg. Med. Chem. 2016, 24, 4812–4825.

43. Vanommeslaeghe, K.; Hatcher, E.; Acharya, C.; Kundu, S.; Zhong, S.; Shim, J.; Darian, E.; Guvench, O.; Lopes, P.; Vorobyov, I., CHARMM general force field: A force field for drug-like molecules compatible with the CHARMM all-atom additive biological force fields. J. Comput. Chem. 2010, 31, 671–690.

44. Vanommeslaeghe, K.; MacKerell Jr, A. D., Automation of the CHARMM General Force Field (CGenFF) I: bond perception and atom typing. J. Chem. Inf. Model. 2012, 52, 3144–3154.

45. Vanommeslaeghe, K.; Raman, E. P.; MacKerell Jr, A. D., Automation of the CHARMM General Force Field (CGenFF) II: assignment of bonded parameters and partial atomic charges. J. Chem. Inf. Model. 2012, 52, 3155–3168.

46. Yu, W.; He, X.; Vanommeslaeghe, K.; MacKerell Jr, A. D., Extension of the CHARMM general force field to sulfonyl-containing compounds and its utility in biomolecular simulations. J. Comput. Chem. 2012, 33, 2451–2468.

47. Martínez, L.; Andrade, R.; Birgin, E. G.; Martínez, J. M., PACKMOL: a package for building initial configurations for molecular dynamics simulations. J. Comput. Chem. 2009, 30, 2157–2164.

48. Humphrey, W.; Dalke, A.; Schulten, K., VMD: visual molecular dynamics. J. Mol. Graph. 1996, 14, 33–38.

49. Gowers, R. J.; Linke, M.; Barnoud, J.; Reddy, T. J. E.; Melo, M. N.; Seyler, S. L.; Domanski, J.; Dotson, D. L.; Buchoux, S.; Kenney, I. M. MDAnalysis: a Python package for the rapid analysis of molecular dynamics simulations. Proceedings of the 15th Python in Science Conference, Austin, Texas, United States, July, 2016.

50. Michaud-Agrawal, N.; Denning, E. J.; Woolf, T. B.; Beckstein, O., MDAnalysis: a toolkit for the analysis of molecular dynamics simulations. J. Comput. Chem. 2011, 32, 2319–2327.

