## Supplemental Figure S1-S6 for "Optical Control of Neuronal Activities with Photoswitchable Nanovesicles"

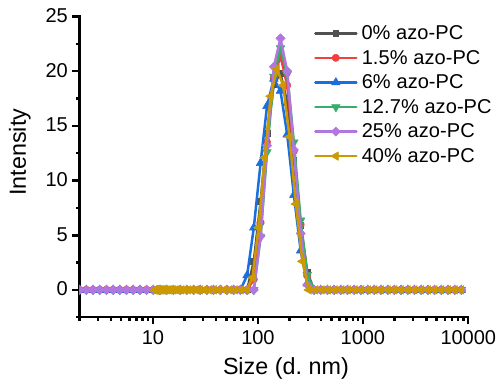

Figure S1. The size distribution of azosomes with different percentages of azo-PC.

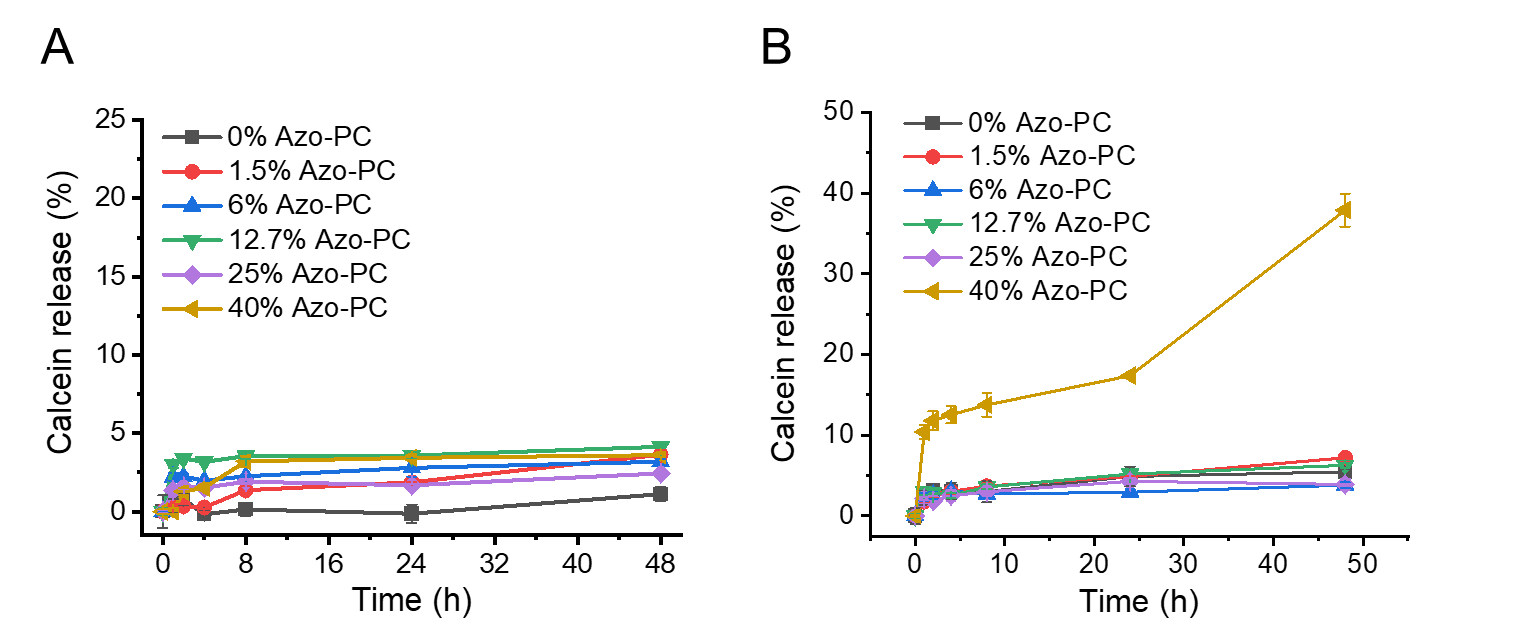

Figure S2. The calcein leakage from azosomes with different percentages of azo-PC at (A) 22^o^C and (B) 37 ^o^C in 0.01 M PBS. Data are expressed as Mean ± S.D.

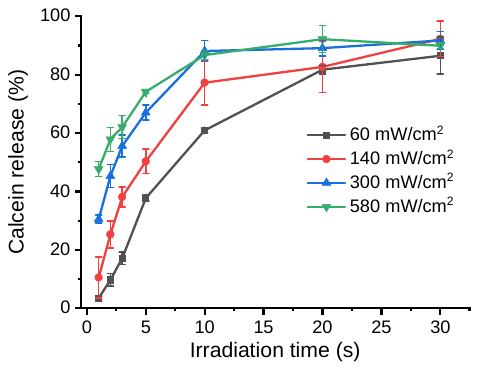

Figure S3. Calcein relelase efficiency at different light densities and durations upon 405 nm light irradiation (azo-PC: 12%). Data are expressed as Mean ± S.D.

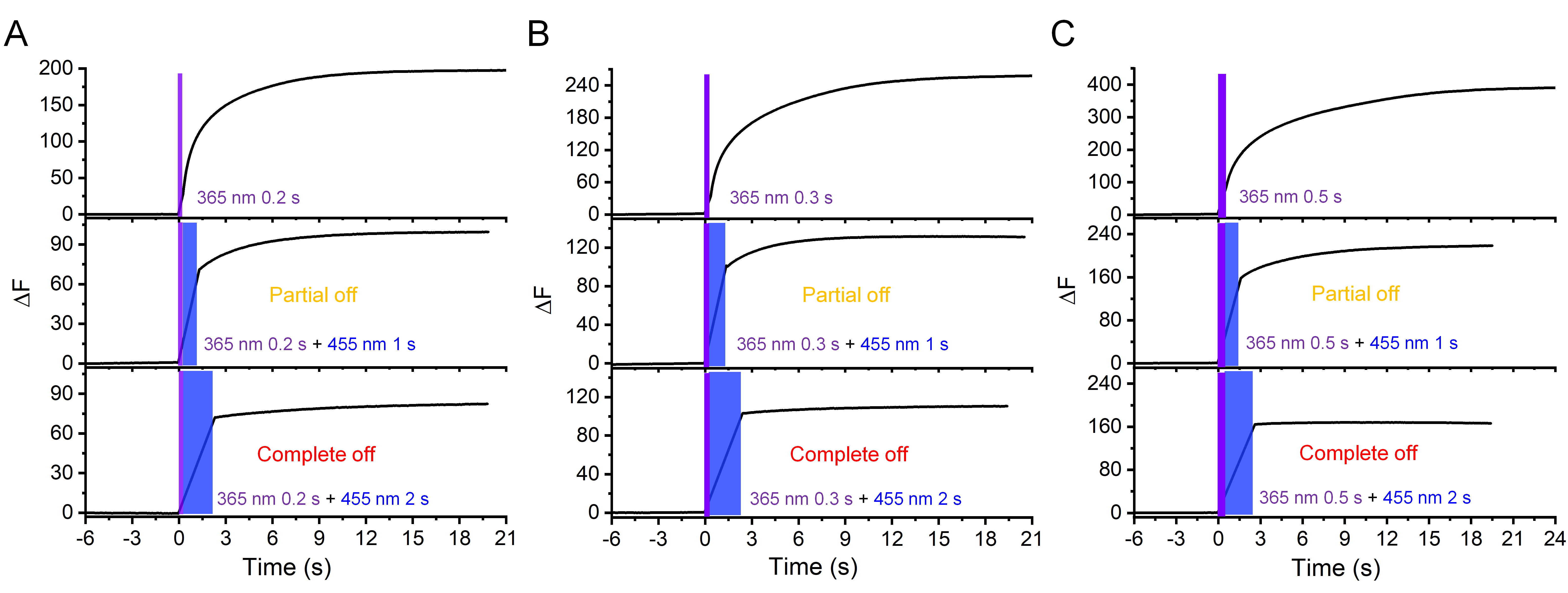

Figure S4. Real-time fluorescence intensity of calcein-azosome over time under the single irradiation of 365 nm and 455 nm light (40 mW/cm^2^). The durations of 365 nm irradiation were 0.2s, 0.3s and 0.5s in (A), (B) and (C), respectively.

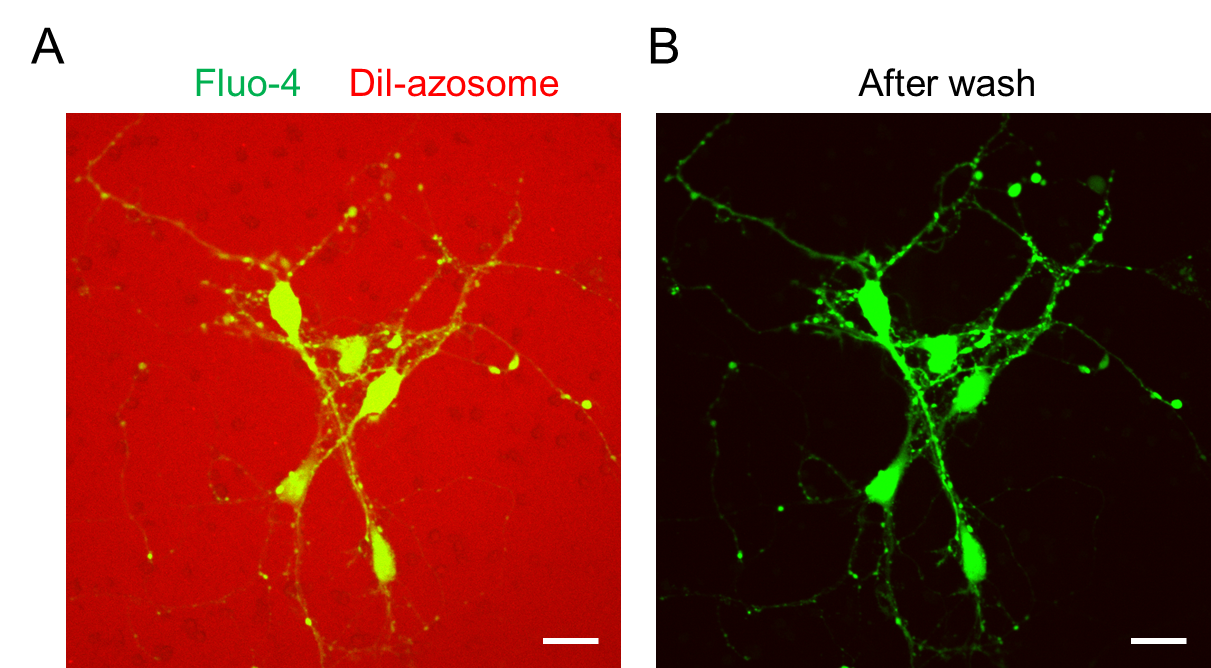

Figure S5. (A) Confocal images of live mouse primary neurons incubated with Dil-azosome (red) for 1 h. Neurons were labelled by fluo-4 (green). The concentration of azosome was 10-fold higher than that in the neuromodulation experiments. (B) Confocal images of live mouse primary neurons after wash. Nearly no fluorescence of Dil-azosome was observed after wash, suggesting that azosomes were not endocytosed by the neurons during the incubation. Scale bar: 20 µm.

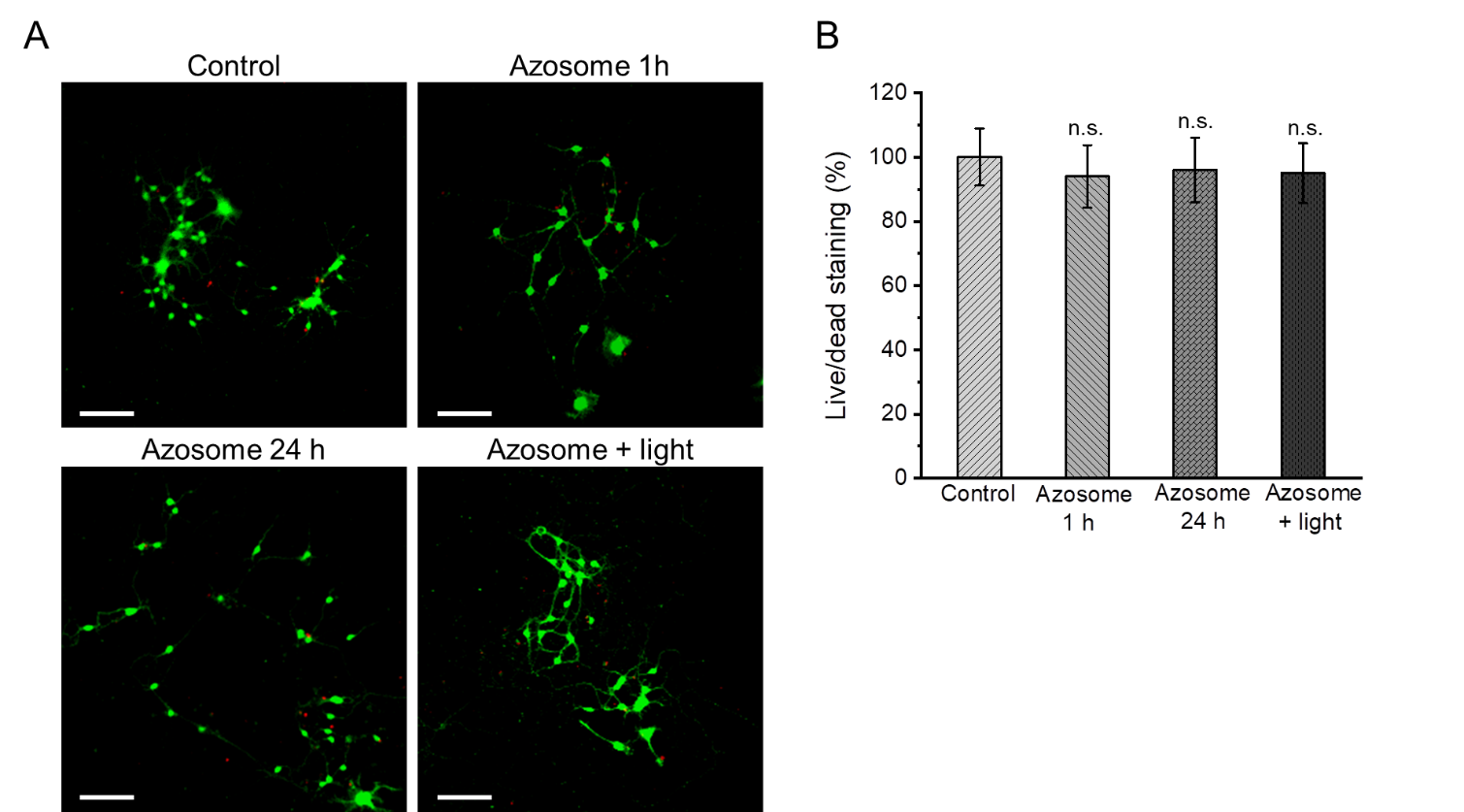

Figure S6. *In vitro* cytotoxicity of azosome. (A) Confocal images of mouse primary neurons stained by calcein (green) and PI (red). Scale bar: 100 µm. (B) The fluorescence intensity ratio of calcein and PI measured by the plate reader after different treatments. No significant difference was found between the control group and azosome-treated groups by two-sample Student’s t-test.
